# Combining predictive coding with neural oscillations optimizes on-line speech processing

**DOI:** 10.1101/477588

**Authors:** Sevada Hovsepyan, Itsaso Olasagasti, Anne-Lise Giraud

## Abstract

Speech comprehension requires segmenting continuous speech to connect it *on-line* with discrete linguistic neural representations. This process relies on theta-gamma oscillation coupling, which tracks syllables and encodes them in decipherable neural activity. Speech comprehension also strongly depends on contextual cues predicting speech structure and content. To explore the effects of theta-gamma coupling on bottom-up/top-down dynamics during on-line speech perception, we designed a generative model that can recognize syllable sequences in continuous speech. The model uses theta oscillations to detect syllable onsets and align both gamma-rate encoding activity with syllable boundaries and predictions with speech input. We observed that the model performed best when theta oscillations were used to align gamma units with input syllables, i.e. when bidirectional information flows were coordinated, and internal timing knowledge was exploited. This work demonstrates that notions of predictive coding and neural oscillations can usefully be brought together to account for dynamic *on-line* sensory processing.

## INTRODUCTION

Neural oscillations are involved in many different cognitive operations (Buzsáki and Draguhn, 2004; Lakatos *et al.*, 2008; Wang, 2010), and considering their cross-frequency coupling permits to even more closely approach their function, e.g. perception, memory, attention etc. (Hyafil, Giraud, *et al.*, 2015). In the domain of natural speech recognition, an important role has been assigned to the coupling of *theta* and *gamma* oscillations (Canolty *et al.*, 2006; Ghitza, 2011; Giraud and Poeppel, 2012), as it permits to hierarchically coordinate the encoding of phonemes within syllables, without prior knowledge of their duration and temporal occurrence, i.e. in a purely bottom-up on-line way (Hyafil, Fontolan, *et al.*, 2015). Natural speech recognition also strongly relies on contextual cues to anticipate what is going to be said next, and when (Rimmele *et al.*, 2018). Recent studies underline the importance of top-down predictive mechanisms during continuous speech perception and relate them to another range of oscillatory activity, the low-*beta* band (Ghitza, 2011; Fontolan *et al.*, 2014; Park *et al.*, 2015; Lewis *et al.*, 2016; Sedley *et al.*, 2016; Pefkou *et al.*, 2017). Predictive coding (Rao and Ballard, 1999; Friston and Kiebel, 2009; Bastos *et al.*, 2012) offers a theory of brain function in the tradition of Analysis-by-Synthesis (Liberman *et al.*, 1967; Norris and McQueen, 2008; Moulin-Frier *et al.*, 2015) and the Bayesian Brain (Knill and Pouget, 2004) hypothesis, which are invoked as critical in speech processing (Poeppel, Idsardi and Van Wassenhove, 2008).

Bottom-up and top-down approaches of speech processing both find support in modeling studies. A neurocomputational model involving the coupling of realistic *theta* and *gamma* excitatory/inhibitory networks was able to pre-process speech in such a way that it could then be correctly decoded by machine learning (Hyafil, Fontolan, *et al.*, 2015). This model aimed at understanding the computational potential of realistic oscillatory neural processes rather than simply fitting existing data. A radically different model, solely based on predictive coding, was able faithfully recognize isolated speech items (such as words, or full sentences when considered as a single speech item) (Yildiz, von Kriegstein and Kiebel, 2013). Although both approaches intended to describe speech perception, one model focused on the *on-line* parsing aspect of speech processing, and the other on recognizing isolated speech segments (no parsing needed). Combining the physiological notion of neural oscillations with the cognitive notion of predictions is appealing as it could broaden the capacity, improve performance, and enhance the biological realism of neurocomputational models of speech processing. More generally and interestingly, such an attempt offers the opportunity to explore the possible articulations between two equally important neuroscientific levels of description, computational/algorithmic for analysis-by-synthesis and algorithmic/implementational for neural oscillations (Marr and Poggio, 1976).

In this study, we addressed whether a predictive coding speech recognition model could benefit from neural oscillation processes. We designed a neurocomputational model based on the predictive coding framework in which we included theta and gamma oscillatory functions. The specific goal of the model was to parse and identify *on-line* syllables from natural sentences. We examined the possible mechanisms by which theta oscillations can interact with bottom-up and top-down information flows and assessed the effects of this interaction on the efficacy of the syllable decoding process. We show that *on-line* speech recognition works best when syllable onset information provided by theta oscillations is combined with endogenous knowledge about syllable duration, and more broadly when continuous predictive processes are informed by dynamic oscillation-based cues.

## RESULTS

### Model architecture and performance

Our goal was to assess the role of temporal information/cues in the extraction of individual syllables from a continuous speech signal (segmentation). We hypothesized that internal generative models including temporal predictions should benefit from such cues. To address this hypothesis and to account for recurrent processes occurring during speech recognition (Wacongne *et al.*, 2011; Gagnepain, Henson and Davis, 2012; Fontolan *et al.*, 2014; Lewis and Bastiaansen, 2015), we used a continuous predictive coding model (described in Methods). Our model explicitly separates “what” from “when”, with “what” referring to the identity of a syllable and its spectral representation (a non-temporal sequence of spectral vectors), and “when” to the prediction of the timing and duration of syllables as implemented through periodic/oscillatory processes (Arnal, 2012; Arnal, Doelling and Poeppel, 2015). “When” predictions took two forms: statistical (not syllable specific) in a theta-module, and syllable-specific in a parallel spectrotemporal-module (Figure 1).

**Figure 1.**
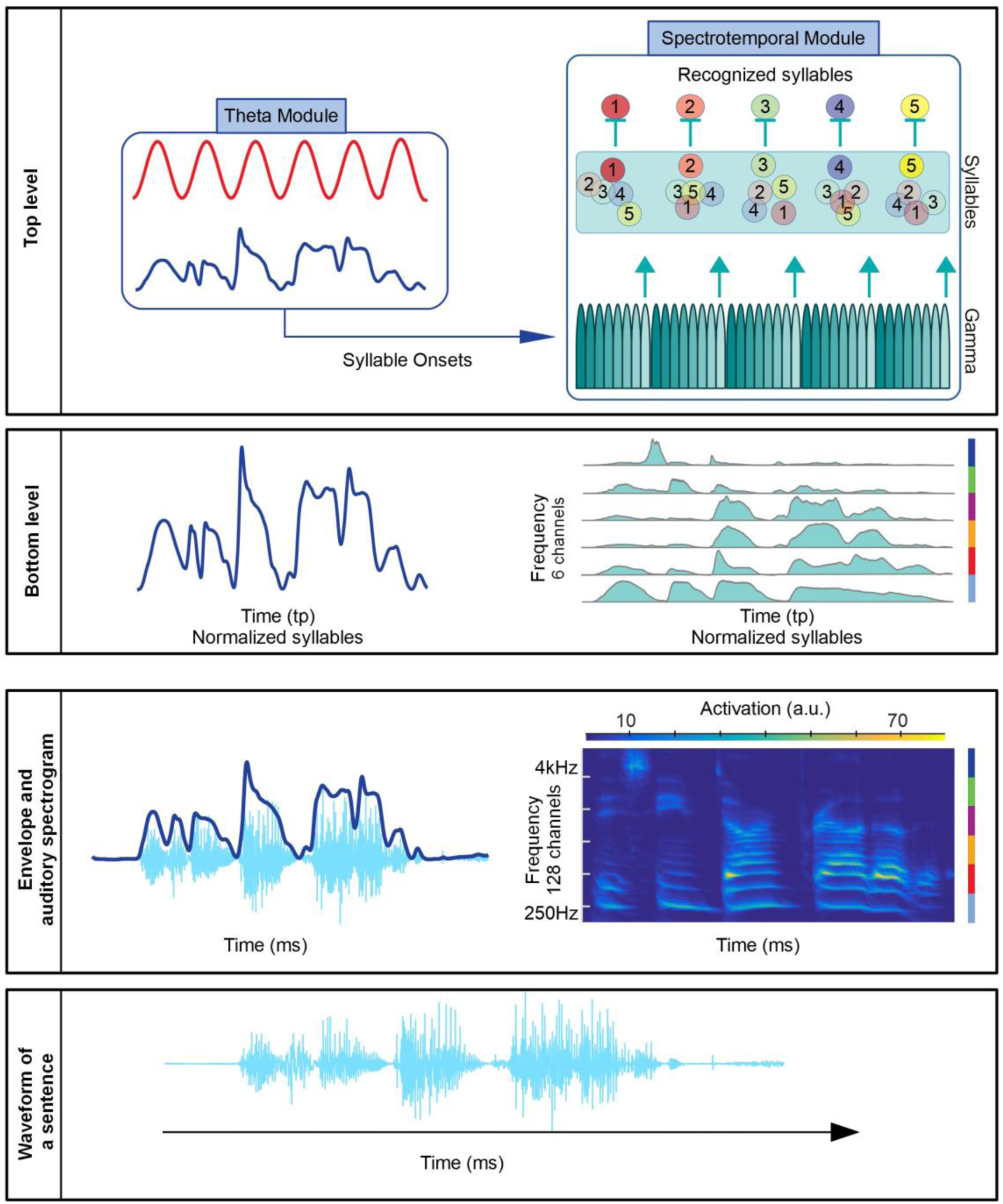
Model of natural sentence processing: syllable parsing and identification. The bottom two rows show the speech waveform and the corresponding envelope and full auditory spectrogram (Chi, Ru and Shamma, 2005). The envelope and a reduced auditory spectrogram serve as input to the model, whose two main levels are represented in the top two panels. The bottom level of the generative model encodes the envelope (left) and the dynamics of the reduced 6 frequency channels (right). The reduced auditory spectrogram was obtained by splitting the original 128 channels into 6 bands; the extent of each band is indicated by a different color in the stacked vertical lines to the right of the spectrogram. The averages across frequencies within each band define the reduced channels appearing on the bottom level of the model (the colors of the stacked vertical lines indicate the corresponding frequency band in the full spectrogram). The top level of the model includes a theta module, and a spectrotemporal module with gamma and syllable units. The theta module contains a theta oscillation and is fed with the envelope; it extracts candidate syllable onsets from local minima in the envelope and uses a theta oscillation to parse them. It sends the resulting signal to reset gamma units. The 8 gamma units operate at the gamma scale and provide processing windows for the syllable encoding process. They activate sequentially and periodically at syllabic rhythm. The model uses the last (8^th^) unit in the cycle to “predict” the end of the syllable and reset syllable units to a common value, such that evidence accumulation can start anew upon arrival of a new syllable (upward arrows). The gamma units also deploy in time the learned spectrotemporal patterns associated with each syllable unit. During inference, the model changes the activation level of each syllable unit, which represents the model’s estimated probability that the associated syllable is generating the current sensory input. The unit with the highest average activation level within the temporal window corresponding to a syllable in the input sentence is considered the recognized syllable.

The model works by inverting an internal model that generates sensory input from internal representations about the causes of that input. Sensory input corresponds to the speech envelope and a 6-channel auditory spectrogram of a full natural sentence (bottom rows of Figure 1), which the internal model generates from four elements (depicted in Figure 1): 1/ a theta oscillation, 2/ a speech envelope unit in a *theta*-module, 3/ a pool of syllable units (as many as syllables in the input sentence), and 4/ a bank of eight gamma units in a spectrotemporal-module. Together gamma and syllable units generate top-down predictions of the input spectrogram. Each of the eight gamma units represents a phase in the syllable; they activate sequentially and the whole activation sequence repeats at syllabic rhythm. Each syllable unit is hence associated with a sequence of eight vectors (1 per gamma unit) with 6 components each (one per frequency channel) (see Figure S1). The auditory spectrogram of a single syllable is therefore generated by the dynamic activation of the corresponding syllable unit over the entire duration of the syllable. As gamma units become sequentially activated they span the 8 vectors that represent the syllable and are deployed over the duration of the syllable (400 timepoints in the simulation, corresponding to 200ms).

Since each gamma unit corresponds to a specific vector in the sequence, it is important that the 1^st^ gamma unit aligns with the beginning of a syllable in the input. To achieve this precise alignment the theta-module detects syllable onsets making use of an envelope unit that tracks the envelope, and a theta oscillation. Syllable onset detection relies on the selection, by the theta oscillation, of local minima in the envelope that are separated by (at least) one theta period. When local minima in the envelope fall within the eligibility window defined by a specific theta phase, they are considered as syllable onsets (Figure 2C). Detected syllable onsets reset gamma units so that the first one, which initiates the whole sequence, is activated at the right time. Once gamma units are temporally aligned with the input, the model updates its estimates about syllable units to minimize the difference between the model’s generated spectrogram and the actual input spectrogram. Syllable units whose spectrogram is consistent with the sensory input increase their activity level, while activity of the others decreases. Successful recognition occurs when this online prediction error driven process leads to elevated activity in a single syllable unit. The syllable unit with the highest activation level within the temporal window corresponding to a syllable in the input sentence is considered the recognized syllable, Figure 2D. The last (8^th^) gamma unit (Figure 2E, teal arrows) signals the last part of the syllable and resets all the syllable units to a common low activation level; thus, evidence accumulation can start anew when the sensory input from the next syllable arrives.

**Figure 2.**
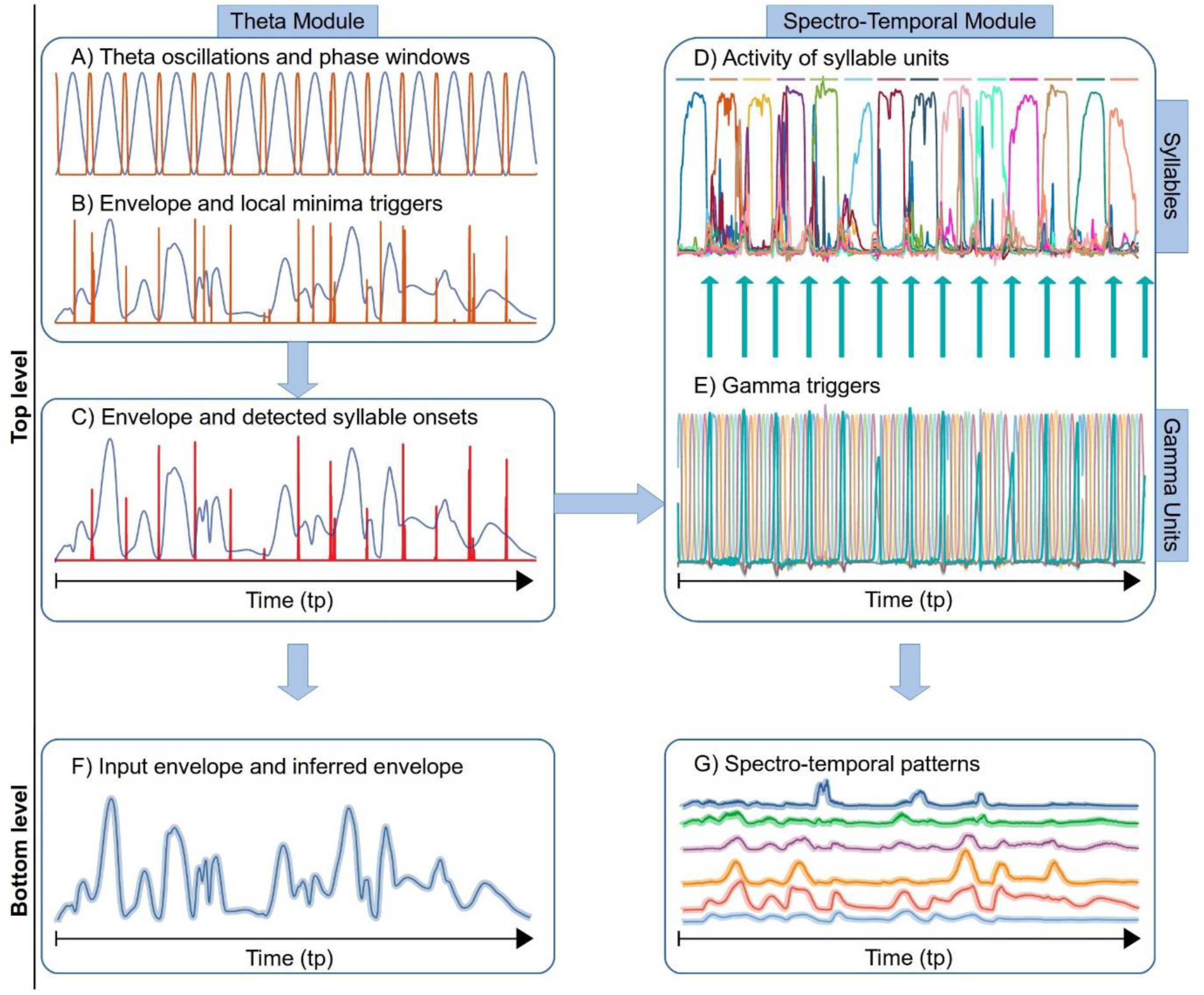
Temporal dynamics of model variables during the inference process. Panels A, B and C on the left illustrate the extraction of theta and local minima triggers, the two alternative syllable onset signals that can be extracted from the speech envelope. Panel A: theta oscillation (in blue) and eligibility windows defined by theta-phase (in red). Panel B: envelope causal state (in blue) and local minima triggers derived from it. The outcome of theta filtering is shown in panel C with envelope in blue and, in red the level of coincidence between the eligibility windows in A (derived from the theta oscillation) and local minima in B (derived from the speech envelope). Red spikes correspond to the model’s estimate of syllable onsets, and define the triggers used to reset gamma units in the full model. The theta module also generates a speech envelope signal (panel F, thick line), which is compared with the input envelope (thin line) at the bottom level. Panels D and E illustrate the reset of gamma and syllable units (causal states in E and D, respectively). Syllable onset information carried by theta triggers resets the dynamics of gamma units and the last gamma unit (thick teal coloured unit, *T_int_*) is used to reset the dynamics of syllable units. Causal states corresponding to gamma and syllable units generate the sound spectrogram (panel G, colour codes the 6 frequency channels) as a weighted sum of syllable specific patterns fed into a Hopfield network (detailed in equations 18 and 19, in Methods). The generated envelope (2F, thick line) and amplitude modulations of the 6 frequency channels (2G, thick lines) are then compared with the envelope (2F, thin line) and auditory spectrogram (2G, thin lines) of the input sentence; mismatch between predicted dynamics and input (prediction errors) propagates back through the hierarchy and updates internal estimates of the hidden and causal states in the theta and spectrotemporal modules. The sentence used for these illustrations was “She had your dark suit in greasy wash water all year”.

In the model, theta and gamma oscillations are used for syllable parsing and spectrotemporal pattern encoding at phonemic scale, respectively. The theta module implements an explicit oscillator at theta frequency, deploys temporal predictions about syllable onset and results in syllable parsing. It represents endogenous knowledge about average syllable duration and is not syllable specific. The spectrotemporal module is based on a theta phase code at phonemic scale for the deployment of the syllable spectrogram, each gamma unit representing a given phase within the overall syllable-driven theta. This module contains predictions about spectral features and expected duration of each possible syllable. These predictions are deployed in a nested way by having gamma units repeat at a theta rhythm (Figure 1, top-level). In summary, syllable specificity is implemented through syllable units in a pool of potential syllables. Syllable units express the model’s confidence that the corresponding syllable is the cause of the current sensory input; their causal states take values from 0 to 1. The model can therefore make any number of syllable-specific predictions based on the activation of the syllable units. In the present model, they predict a spectral pattern (a sequence of spectral vectors) and implicitly their own duration, which corresponds to a fixed sequence of gamma units, where each one represents a particular phase within a syllable specific “theta” period (a theta-related principle different from the one used within the theta-module).

The model was presented with 30 spoken English sentences corresponding to 10 different sentences for 3 different speakers (Table S1). Sentences were presented one at a time and the syllable pool contained only the syllables present in the input sentence. Even though a biological theta oscillator can deal with variable syllable rates (Hyafil, Fontolan, *et al.*, 2015; Pefkou *et al.*, 2017), we normalized syllable length to have standardized spectrotemporal representation and same number of gamma units for each syllable. After normalization all syllables had the same duration. However, naturally occurring gaps of different durations broke perfect rhythmicity. Figure 2 illustrates the dynamics of the model’s first and second levels during inference for a sample sentence. Model performance, quantified by the median percentage of correctly identified syllables was 90.2%. For the sentence in Figure 2, all syllables were correctly identified. After each reset of syllable units at the end of the previous syllable, all units started with the same low activation and prediction errors increased the activity of the correct unit (Figure 2D).

### Resets and theta oscillation

The model presented above includes a physiologically motivated theta oscillation, which uses local minima in the envelope to provide syllable onsets, to temporally align internally generated predictions with the input. It also relies on the reset of accumulated evidence by silencing syllable units at syllable boundaries. In a next step, we sought to assess the importance of these two processes for proper online syllable parsing and identification. We also wanted to compare the two sources of temporal information available, the endogenous knowledge represented by gamma and syllable units, and the external information carried by the combination of envelope and theta oscillation. To this end, we run simulations for five different model architectures (Figure 3, top panel), where the main components were the same, but the information used to organize (reset) the activity of gamma and syllable units differed (Table 1, in Methods). Model 3E is the model described in the previous section and depicted in Figure 1. Variants included models with 1/ no reset of syllable units (no reset of accumulated evidence, models 3A and 3B), 2/ reset of gamma units by candidate syllable onsets provided directly by local envelope minima (3A and 3C), or 3/ by theta parsed local minima (3B, 3D and 3E), and 4/ reset of syllable units (accumulated evidence) by theta parsed local minima (stimulus driven duration information, 3D) or 5/ by the last gamma unit (internal expectations of syllable duration, 3E).

**Table 1.** Triggers for each model configuration

|  | Gamma units | Syllable units |
| --- | --- | --- |
| Model A | $T_\gamma = T_{lm}$ | $T_\omega = 0$ |
| Model B | $T_\gamma = T_\theta$ | $T_\omega = 0$ |
| Model C | $T_\gamma = T_{lm}$ | $T_\omega = T_{int}$ |
| Model D | $T_\gamma = T_\theta$ | $T_\omega = T_\theta$ |
| Model E | $T_\gamma = T_\theta$ | $T_\omega = T_{int}$ |

**Figure 3.**
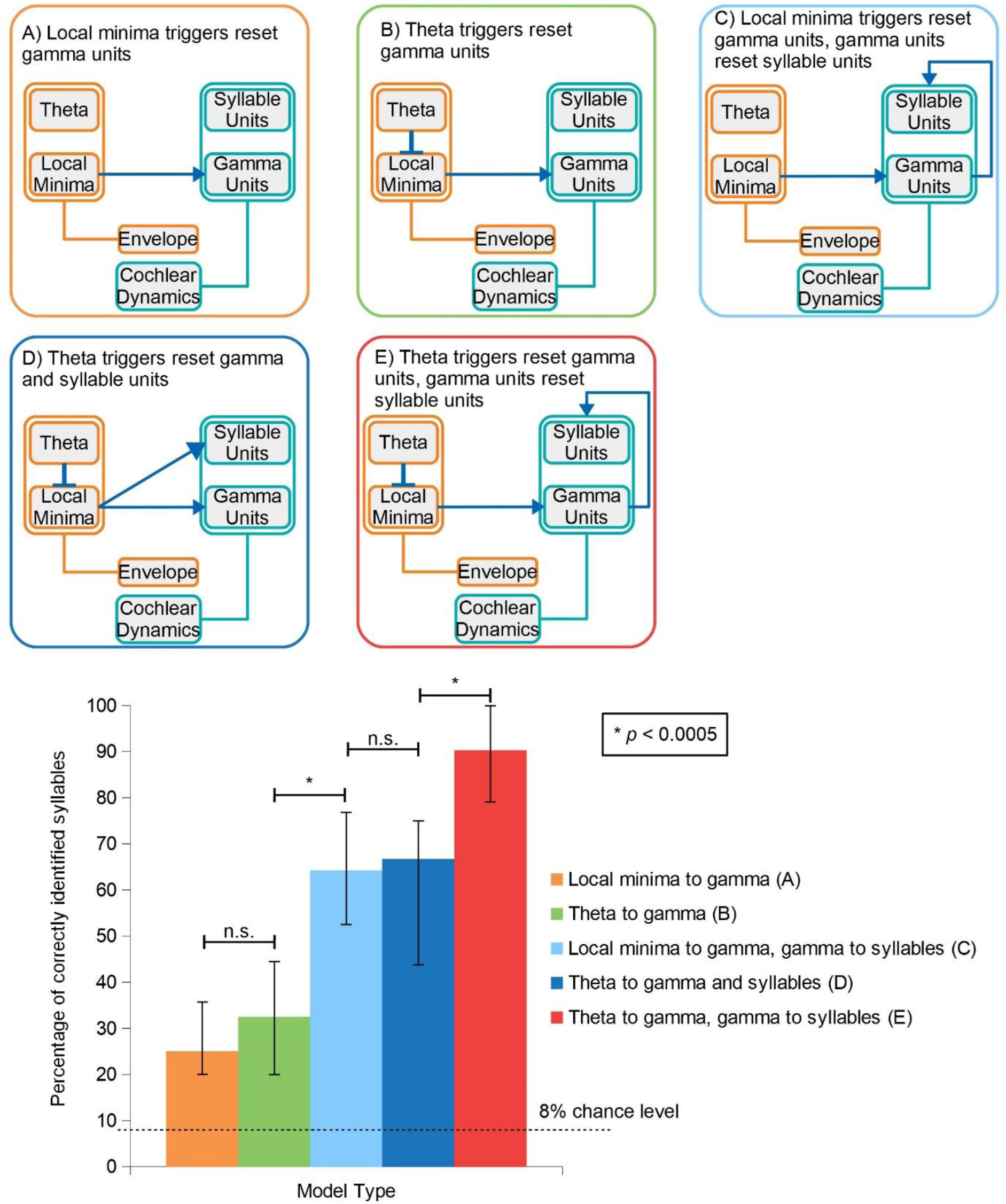
Model variants and their performance. To assess the relative contribution of each source of temporal information, we tested 5 configurations of the model (top panel). The bar plot represents the median performance of each model configuration on the 30 sentences, with error bars showing the 25% and 75% percentiles across the 30 sentences; the dashed horizontal line represents chance level. Pairwise comparisons were performed using the Wilcoxon signed rank test with false discovery rate to correct for multiple comparisons.

Figure 3 (bottom panel) shows the median performance for each architecture. Although all model variants performed well above chance level (around 8%), there were significant performance differences across them (p-values for pairwise comparisons are presented in Table S2). Performance was significantly lower in models A and B versus C, D, and E, indicating that syllable units reset is a crucial factor for accurate speech encoding, as it cancels out accumulated evidence about previous syllable before the processing of a new syllable starts. Furthermore, when we compared models A and C versus B and E, we found that using theta triggers to reset gamma units (B and E) rather than local envelope minima triggers (A and C) also improved model’s performance. Interestingly, the increase in performance due to theta triggers is small (and non-significant) when the reset of syllable units was disabled (A vs B). In comparison, when syllable units were reset, the performance gain caused by theta triggers was around 30% and statistically significant (C vs E). The conclusion of these simulations is that the model performs best when gamma units are reset by theta oscillations, and when syllable units are endogenously reset after completion of each gamma-units sequence. This means both stimulus-driven information filtered by theta oscillations and reset of evidence accumulation are equally important.

However, two different mechanisms can reset syllable units. In model variant D, syllable units are reset by syllable onsets as detected by the theta oscillation, whereas in model variant E syllable units are endogenously reset by the last gamma unit. Our simulations indicated that model variant E performed better than D, meaning that performance benefited from the combination of external, envelope driven syllable onsets, with endogenously generated syllable duration information. Figure S2 illustrates how each trigger impacts the internal dynamics of syllable and gamma units for different model architectures. The figure clearly shows that syllable onsets detected by the theta oscillation contribute to temporally organize gamma units, whereas explicit reset of syllable units results in clear peaks within syllable boundaries, hence in better recognition. Finally, the model variant depicted in Figure 3E, which combines theta detected syllable onsets and internal expectation about syllable durations to reset gamma and syllable units, respectively, yielded the least distorted gamma sequence activity (no breakdown of their orderly sequential activation) and clear winners within each syllable boundaries (Figure S2E).

Overall, these results show that two processes positively influence online speech segmentation into distinct recognisable syllables. The first one pertains to the implementation of the coupling of theta/gamma processing levels: gamma units must be reset by syllable onset as detected by theta oscillations, while evidence accumulation by syllable units must be reset by gamma units upon syllable completion. The second one pertains to the ability of the model to combine stimulus driven information (theta triggered onsets) and endogenous temporal information (gamma-related syllable-specific duration).

## DISCUSSION

We designed a hierarchical on-line speech recognition model, composed of a first level that receives as input the envelope and time-frequency decomposition of natural English sentences, and a second level that involves a theta-oscillation and a module containing a sound spectrogram. The role of the theta module is to parse the input sentence into syllable-like segments by picking local minima in the speech envelope that correspond to syllable onsets. This information is then used to reset gamma-rate activity in the spectrotemporal module, ensuring that the latter aligns with syllable timing in the input. Gamma and syllable units dynamically generate the spectral content of syllables, and each syllable is encoded by the sequential activation of a bank of eight 25ms integration gamma units (Roß *et al.*, 2000; Lakatos, 2005). Model simulations suggest that on-line syllable parsing and recognition requires an explicit reset of syllable units before the syllable starts or after it ends. The model’s performance is higher when this explicit reset is based on expected syllable duration rather than only on syllable onset information extracted by theta oscillations. However, the best model performance is obtained when 1) theta oscillations reset gamma units and 2) gamma units reset syllable units, a model architecture where theta oscillations produce an optimal alignment of top-down and bottom-up information flows. In summary, our simulation results suggest that *on-line* speech recognition works best when syllable onset information delivered by theta oscillations is used in combination with internal knowledge about syllable duration, and more generally when dynamic oscillation-based cues interact with continuous predictive processes.

The proposed model was exapted from a speech recognition model (Yildiz, von Kriegstein and Kiebel, 2013), inspired by birdsong generation (Yildiz and Kiebel, 2011). While most simulations described in the 2013 article were performed on monosyllable words (digits), our proposed model extends the approach to natural continuous speech sequences made up of subunits (syllables in our case) whose order is not known a priori, an important step toward neurobiological realism of neurocomputational models of language processing. Indeed, speech is made of linguistic building blocks permitting (quasi)infinite combinatorial possibilities. We capture this combinatorial freedom by assuming that syllables can appear in any order, i.e. have an equal probability, which is not entirely true for natural speech. This place our model in a more challenging situation than when facing real speech statistics, which emphasizes our main results that the precise coordination of bottom-up and top-down information flow is critical for recognition, and that neural oscillations optimize this coordination by signalling syllable boundaries. In our model, the input is a sentence that can vary in duration from 6 to 23 syllables separated by gaps of different durations. The first and essential step the model has to achieve is therefore to retrieve correct syllable boundaries. This syllabification issue is non-trivial and is classically dealt with by *off-line* methods, e.g. Mermelstein algorithm (Mermelstein, 1975). Yet, a theta-range natural oscillator can achieve accurate *on-line* syllabification (Hyafil, Fontolan, *et al.*, 2015). This is possible when the oscillator is *weak*, meaning that its intrinsic frequency can adapt, within limits, to that of an external stimulus. The theta oscillator in that case was built from reciprocal connections between excitatory and inhibitory leaky integrate-and-fire neurons. For the current model, we used a simplified version of this network that enabled online syllable onset detection based on envelope tracking. Note that a more biophysical theta oscillator flexibly driving integration windows in the gamma-unit bank would allow us to deal better with syllables of variable length, as they occur continuous natural speech. For the sake of parsimony, this was not implemented in the present work.

A critical aspect of the model is the coupling between theta and gamma modules. Many experimental studies show that neural theta and gamma activity interact (Lakatos *et al.*, 2008; Luo and Poeppel, 2012; Lisman and Jensen, 2013; Hyafil, Giraud, *et al.*, 2015; Lam *et al.*, 2016), and most likely that theta organizes gamma activity to align neural encoding timing with stimulus timing to preserve the hierarchy between phonemes and syllables (Giraud and Poeppel, 2012; Martin and Doumas, 2017). Although this is not the only possible option (Kösem *et al.*, 2016), in our variant models B, D and E (Figure 3) we adopt the most straightforward view (Lakatos, 2005; Ghitza, 2011) that the slow oscillation controls the fast one, and implemented it via a reset of gamma units by theta triggers. The observation that theta triggers always performed better than local minima triggers (A and C versus B, D and E), shows that the theta module successfully filters out irrelevant acoustic troughs in the envelope, which is a challenge for envelope-based syllable boundary detection (Villing, Ward and Timoney, 2006). Comparing the performance of models A vs B and C vs E, further suggests that the temporal organization of gamma units by theta triggers results in better coordination of top-down and bottom-up information flows, than when all local minima are taken into account. The model also contains a gamma-theta interaction within the spectrotemporal module, where gamma units by themselves build their own theta rhythm. This implicit coupling provides an additional syllable parsing opportunity.

Although important, the reset of gamma units by the theta module was not sufficient to ensure good performance. The latter remained modest, around 30% (i.e. 20% above chance level), in all models that did not further include explicit syllable units reset. If a syllable is recognized, the corresponding syllable unit is higher than the others and affects the processing of the next syllable. By incorporating a reset of the accumulated evidence about the previous syllable, performance increased by up to 50% (models A and B versus C, D and E, Figure 3). However, only model variant 3E, which combines envelope driven temporal information (theta triggers) with endogenous syllable duration information (gamma triggers) performed significantly higher than model variant D that only relies on theta triggers. There are two possible reasons for this difference. First, as theta triggers reset syllable units at the beginning of a syllable, and gamma triggers at the end, there could be a reset position effect on the model's performance. Second, the theta module can occasionally fail to detect some syllable onsets, thus failing to reset syllable units and preventing correct identification. To test this hypothesis, we re-ran simulations on these two architectures (D and E), with a modification. Instead of leaving the model detect syllable onsets from the envelope, we explicitly provided syllable onset information to the model. Figure 4 shows that the modified model correctly identifies all the syllables for both variants (theta triggers with or without gamma triggers). In sum, when all syllable onsets are ideally detected the model is always able to recognize syllables. Arguably in realistic listening conditions, when not all syllable onsets are detected (because of noise, cocktail-party situations etc), the brain might benefit from endogenous clocking mechanism - implemented here as a *fixed* gamma units sequence. Our results hence suggest that some processing redundancy might be necessary to cope with the various challenges posed by natural speech on-line recognition, and that on-line speech recognition heavily relies on a trade-off between adaptability to speech rate via flexible theta oscillations, and relatively fixed encoding rate via endogenous gamma. This observation concurs with our previous experimental findings that theta but not gamma oscillations flexibly track speech rate (Pefkou *et al.*, 2017).

**Figure 4.**
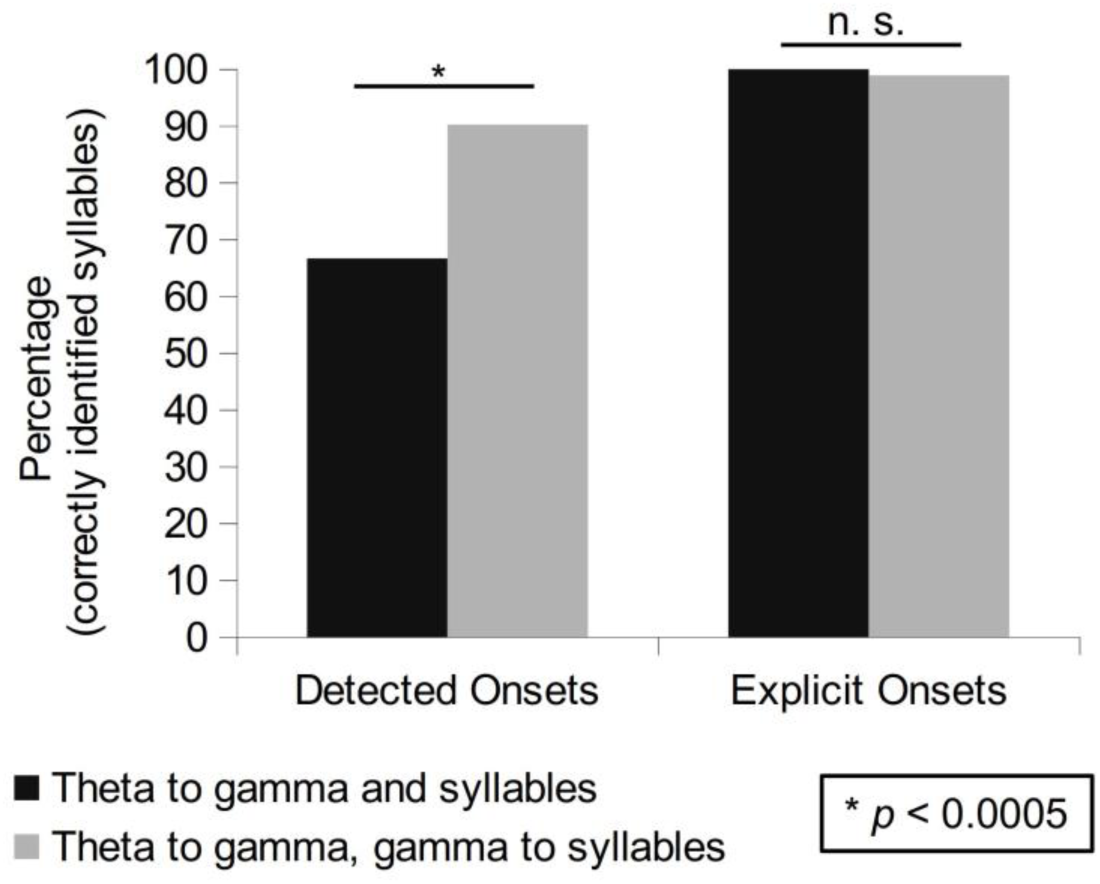
Explicit versus detected onsets. Performance for configurations D and E depending on whether syllable onsets are detected by the theta-module (detected onsets) or when true syllable onsets are explicitly provided to the model (explicit onsets; ideal onset detection condition). For explicit (true) onsets, both model versions show very high performance, therefore whether the triggers to syllable units are provided at syllable onset (configuration D) or end (configuration E) is not the key element for the performance difference. As theta-onset detection is not perfect, especially when there is a big gap (more than 50-60 tp) between syllables, the configuration that relies on the endogenous syllable duration information performs better.

Although the notion of top-down control is constitutively present in our predicting coding implementation, our model still lacks the notion that top-down processes occur preferentially at low-beta rate, as recently demonstrated (Bastos *et al.*, 2012; Fontolan *et al.*, 2014; Lewis *et al.*, 2016; Sedley *et al.*, 2016). The predictive coding model we used here works in a continuous inferential mode, which is discretized only by virtue of syllable timing. Yet, it seems that gamma bottom-up activity is modulated at low-beta rate (Fontolan *et al.*, 2014; Bouton *et al.*, 2018), which could offer top-down integration constants that are intermediate between syllables and gamma phonemic-range chunks, and whose advantage could be to smooth the decoding process by providing sequential priors at intermediate scale between phonemes and syllables. Alternatively, beta top-down rhythm could also be related to *expected precision*, thus encoding second order statistics (Friston and Kiebel, 2009). Expected precision weighs bottom-up prediction errors, hypothesized to work at gamma, and could control their impact on the evidence integration process. When the sensory input corresponding to a new syllable arrives, the large prediction error could decrease the estimated confidence in top-down prediction and boost that in bottom-up prediction error. If the relative weight of bottom-up and top-down information is carried by low beta activity, we would then expect an alternation with a theta rhythm, a finding that was experimentally observed (Fontolan *et al.*, 2014). An important generalization for the model would thus consist in adopting a framework that estimates precisions (Friston, Trujillo-Barreto and Daunizeau, 2008; Friston *et al.*, 2010). These proposals remain speculative, and neurocomputational modelling could be one way to address whether the principle of frequency and temporal division of bottom-up and top-down processing is functionally interesting, and whether low-beta rate for top-down flow is optimal or merely a “just so” phenomenon.

Although our goal was not to design a speech processing model that can compete with those used in the domain of automatic speech recognition (Li *et al.*, 2014; Prabhavalkar *et al.*, 2017; Sak *et al.*, 2017), it turns out that the notion of neural oscillations could be relevant for the latter. Hyafil and Cernak (Hyafil and Cernak, 2015) demonstrated that a biophysically plausible theta oscillator which can syllabify speech *on-line* in a flexible manner makes a speech recognition system more resilient to noise and to variable speech rates. It is possible that introducing more oscillatory mechanisms in ASR could further improve performance and resilience to noise. This is possibly the case for using a basic sampling rate in the low-gamma range as implemented here instead of faster ones as commonly used (Hirsch, Hellwig and Dobler, 2001). Likewise, using oscillation-based top-down updating, which could deploy predictions about when events are expected to happen, something that most ASR systems do not yet do (Davis and Scharenborg, 2016).

This theoretical work shows the interest of extending predictive coding approaches to the domain of neural oscillations, which permits i) to emulate a neurally plausible interface with the real world that is able to deal with the continuous nature of biological stimuli and the difficulty to parse them into elements with discrete representational value, and ii) to provide internal orchestration of the information flow that supply possible deficiencies of interfacing mechanisms.

## Supporting information

## ACKNOWLEDGEMENTS

This work was funded by a grant from the Swiss National Science Foundation to ALG (#320030_163040).

## AUTHORS CONTRIBUTION

ALG and IO designed the study, SH carried out the study under IO and ALG guidance, SH, IO and ALG wrote the manuscript.

## METHODS

### Speech Input

We used 30 recorded English sentences from the TIMIT database (Garofolo *et al.*, 1993) for our simulations. The sentences were spoken by 3 different male speakers (10 sentences each). Overall, those 30 sentences include 389 syllables. Input to the model consisted of 1) a time-frequency representation of the sound wave and 2) the speech envelope. We used a biologically inspired model of the auditory periphery (Chi, Ru and Shamma, 2005) to obtain the time-frequency representation. By default, it transforms the auditory signal into 128 logarithmically spaced frequency channels, spanning from 150 Hz up to 7 kHz. Then we normalized the spectrogram so that its values are between 0 and 1. After averaging the activity of neighboring channels, we reduced the number of channels to 6, covering the range of frequencies from 150 Hz to 5 kHz. To obtain the envelope, we applied the Hilbert transform on the waveform of each sentence (smoothed using Matlab’s (MATLAB 2014b, The MathWorks, Inc., Natick, Massachusetts, United States) default “smooth” function and normalized to [0,1] interval). Each recorded sentence was therefore represented by seven information channels, the envelope (*E*(t)) plus 6 frequency bands (*F_f_*(t); f= 1,… 6).

### Syllabification

The model’s goal is to recognize syllables on-line, which requires defining syllables in the input to subsequently assess the model’s performance. The TIMIT corpus provides phonemic boundaries labeled by professional phoneticians. This information was passed to Tsylab2, a program that provides candidate syllable boundaries based on the TIMIT phonemic annotation and on English grammar rules (Fisher, 1996). In the model, we actually considered an alternative definition of the syllable based on the sound envelope. We used the notion that syllable onsets are located next to local minima of the envelope (Figure S3, dashed black lines on panel c) (Villing, Ward and Timoney, 2006). Most (but not all) local minima in the envelope correspond to moments when the speaker finished/started a speech segment such as a syllable or word. Therefore, we realigned the Tsylab2 boundaries to the closest local minimum on the envelope when the difference between Tsylab2 boundaries and local minimum was below 50 ms; otherwise the Tsylab2 boundaries were kept.

#### Temporal normalization

To keep the model as simple as possible, we normalized syllable duration. Every syllable was represented by 400 time points. The mean syllable duration in our sentence set was around 200 ms; that is, each timepoint corresponds on average to about 0.5 ms. Gaps between syllables were not normalized and resulted in 76 gaps with durations from 7 to 237 timepoints (Figure S4). The resulting set of syllables and gaps was concatenated in the same order as they appeared in the original sentence. The input is, therefore, a sequence of normalized syllables with interleaved gaps of varying duration.

### Generative model

We used a predictive coding model to parse and recognize individual syllables from the continuous acoustic waveform of spoken English sentences. The core of the predictive coding framework is a hierarchically structured generative model that represents the internal knowledge about the statistics and structure of the external world. During the inference process, the brain inverts the generative model and tries to infer the hidden causes of the sensory input. To invert the generative model, we used the Dynamic Expectation Maximization algorithm (Friston, Trujillo-Barreto and Daunizeau, 2008), which is based on top-down predictions and bottom-up prediction errors.

We considered a generative model with two hierarchically related levels. At each level *i* in the hierarchy, dynamics are determined by local hidden states (denoted by *x^(i)^*) and causal states from the level above (denoted by *v^(i)^*). At the same time, each level generates causal states that pass information to the level below (*v^(i-1)^*). Hidden states are subject to dynamical equations while causal states are defined by static, generally nonlinear transformations of hidden states and causal states. Schematically Top level (i = 2)

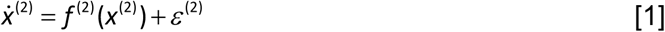

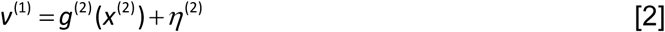

The dynamics at this level are only determined by hidden states *x^(2)^*. *v^(1)^* is the output to the level below. The hidden states at this level include hidden states for a theta oscillator, the speech envelope, syllable units and gamma units (see below for details). ε^*(i)*^ and η^*(i)*^ (*i* = 1, 2) stand for random fluctuations for hidden and causal states respectively (the same notation is used in the next sections); their precision determines how predictions errors are weighted (Friston, Trujillo-Barreto and Daunizeau, 2008). Causal states passing information to the bottom level include causal states for syllable units, gamma units and the envelope.

Bottom level (i = 1)

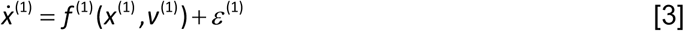

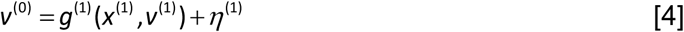

At this level, there are hidden and causal states related to the 6 frequency channels and a causal state for the envelope (which is relayed without modification from the top level).

The output of the bottom level *v^(0)^* is then compared with the input Z(t): a vector containing the envelope and the reduced 6-channel auditory spectrogram.

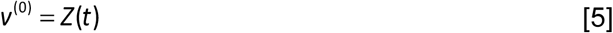

In the following we write the explicit form of these equations. Figure S5 provides a schematic about all the variables used in the model.

### Top Level

The top level has two modules; a theta module with *envelope* and *theta oscillation* units and a spectrotemporal module with *gamma* and *syllable* units.

### Theta Module

The role of this module is twofold. First, it tracks the envelope in the input signal and detects the local minima on it. Second, theta oscillation filters detected local minima that are separated by at least a theta cycle. These are the model’s estimates of syllable onsets.

The theta module tracks and low-pass filters the envelope in the input with the following pair of equations:

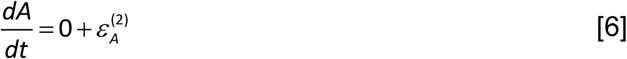

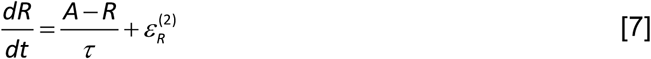

The perfect integrator (equation 6) tracks the amplitude fluctuations of the envelope. During inference, *A* generates an envelope signal that is compared with the input envelope (see Figure S5); precision weighted prediction errors in generalized coordinates drive equation 6 and result in the variable *A* tracking the envelope (Figure 2C). A low-pass filtered version is then provided by equation 7.

Then we wrote equations to extract filtered versions of the first and second order derivatives of the envelope:

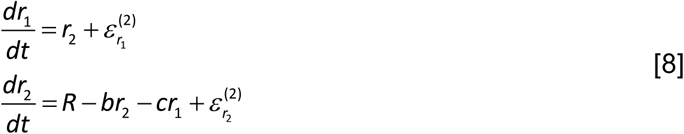

*r_2_* is a filtered version of the first order derivative of *R*.

We apply the same transformation to *r_2_* to obtain *r_2,2_*, a filtered version of the second order derivative of the envelope:

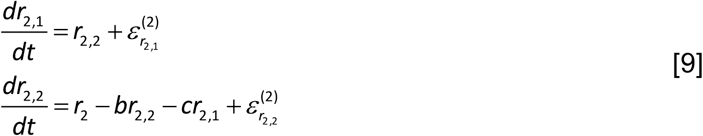

*r_2_* and *r_2,2_* will be used to define the “local minima” in the envelope below (equation 23). Additionally, in the theta module, there is a harmonic oscillator with fixed frequency ‘Ω’:

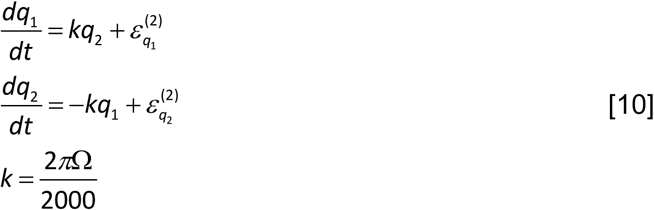

Where 2000 is the sampling rate and Ω = 5 Hz is the syllabic frequency. This ensures that the theta period corresponds to the normalized syllable duration.

### Spectrotemporal module

#### Gamma Units

Gamma units are modeled as a stable heteroclinic channel, which results in their sequential activation (Rabinovich *et al.*, 2006) (for details see (Yildiz and Kiebel, 2011; Yildiz, von Kriegstein and Kiebel, 2013)). The duration of each gamma unit is fixed to 50 timepoints (tp), thus, it operates at a gamma scale (50 tp corresponds to around 25ms). There are 8 gamma units, and the whole sequence of 8 units has the same period as the normalized syllables. In other words, the model has intrinsic information about syllable duration associated with the sequence of the gamma units.

In the model, gamma units provide processing windows for the syllable encoding process. The active unit determines which part of the syllable is encoded at each moment of time. For example, if the first gamma unit is active, then the first 1/8 part of the spectral content of a syllable is encoded, if the second gamma unit is active then the second 1/8 part is encoded and so on.

The mathematical equations are adapted from Yildiz et al (Yildiz, von Kriegstein and Kiebel, 2013).

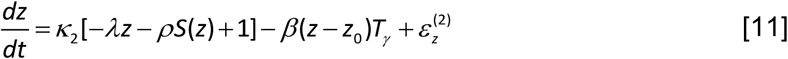

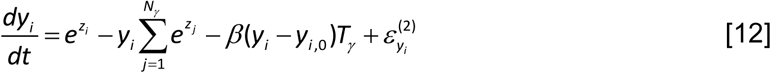

Where

- *i* represents the index of gamma unit and takes values from 1 to N_γ_ = 8
- *z* is a vector of 8 units encoding the amplitude fluctuations of the gamma units, whereas the vector *y* shows the amplitude of the gamma units scaled to [0, 1] interval.
- *z_0_* and *y_0_* represent the reset values of *z* and *y*, corresponding to the state when the first gamma unit is active (the start of the gamma sequence)
- *T_γ_* stands for the trigger that gamma units receive from the theta module (Table 1)
- β = 3 is a scaling factor to amplify triggers
- *S*(z) = *1/(1+e^-z^)* is applied component-wise.
- *ρ_ij_* ≥ 0 is the connectivity matrix, determining the inhibition strength from unit *j* to *i*. The values of the connectivity matrix ρ are defined as follows:

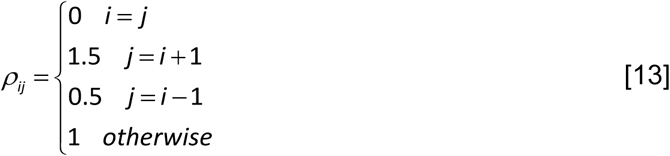

The first term on the right-hand side of both equations is taken from (Yildiz, von Kriegstein and Kiebel, 2013). The value of κ_2_ was set so that the gamma sequence’s duration is 400 tp. We added the 2^nd^ term on the right-hand side of both equations to reset the gamma units whenever a trigger about syllable onset arrives; the triggers ensure that irrespective of the current state of the network, the first unit is activated and the gamma sequence is deployed from the beginning whenever there is a trigger. When the trigger corresponds to a syllable onset, it ensures that the gamma sequence is properly aligned with the input and therefore that the spectrotemporal predictions temporally align with the input. Whenever a syllable starts, the first unit of the gamma sequence should be activated; otherwise the model would encode the spectral content of the wrong part of a syllable.

### Syllable units

The last module of the top level contains the syllable units; they represent evidence that the associated syllables corresponds to the syllable in the input. The number of syllable units varies from sentence to sentence and corresponds to the number of syllables in the input sentence. The equations for syllable units are:

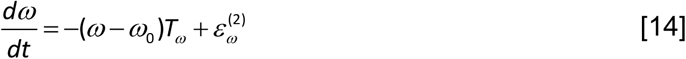

Where T_ω_ corresponds to triggers (Table 1) that reset the activation level of the syllable units. A trigger drives the activity level of all syllable units towards an equal value ω_0_. As we will specify below, triggers originated either from the theta module, signalling estimated syllable onsets, or from the last gamma unit, signalling internal expectations about the end of a syllable. Between triggers syllable units act as evidence accumulators. The activation level of each unit determines its contribution to the generated auditory spectrogram (equations 16 and 19).

The causal states of the second level pass information to the bottom level:

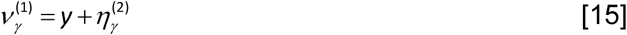

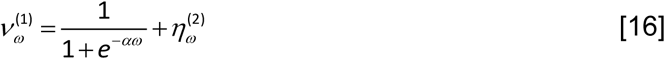

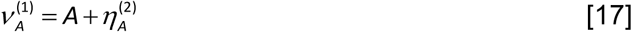

Equation 15 corresponds to the 8 scaled gamma units (equation 12); they activate sequentially and take values between 0 and 1. Equation 16 corresponds to the *N_syl_* syllable units; the corresponding causal states are obtained by applying a component-wise sigmoid function to the syllable hidden states (equation 14) so that they are bounded between 0 and 1. Since all the syllables in the input are present in the memory pool, prediction error (the difference between predicted and actual spectrotemporal patterns at the first level) will be minimized when the causal state of the corresponding syllable unit in the model is driven close to 1 while all others are driven close to 0. Finally, equation 17 sends information about the current estimate of the envelope.

### Bottom level

The bottom level contains variables related to the amplitude fluctuations of the frequency channels as well as the envelope.

The amplitude fluctuations of the frequency channels are modeled with a Hopfield attractor-based neural network (Hopfield, 1982). The following equations were adapted from Yildiz et al (Yildiz, von Kriegstein and Kiebel, 2013).

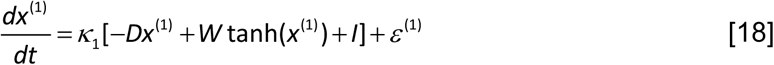

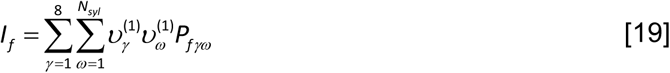

x^(1)^ is a vector with 6 components (one per frequency channel), *D* is a diagonal self-connectivity matrix and *W* is an asymmetric synaptic connectivity matrix; they were designed so that the Hopfield network has a global attractor whose location depends on vector *I* (Yildiz and Kiebel, 2011; Yildiz, von Kriegstein and Kiebel, 2013). In equation 19, *ν*_γ_^(1)^ and *ν*_ω_^(1)^ are the causal states for the gamma and syllable units from the top level (equations 15 and 16). Because of the sequential activation of gamma units during a syllable, *I* represents a vector for each of the gamma units. *P_fγω_* is defined from the spectrotemporal patterns *ST_fγω_* associated with each syllable as follows:

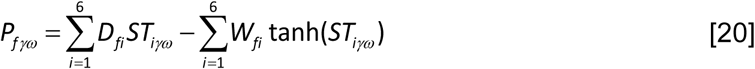

Syllable spectrotemporal patterns *ST_fγω_* were calculated by averaging each of the 6 frequency channels (F_f_(t)) within each of the eight 50tp windows (γ) for each syllable (ω). As the syllables were normalized to 400 tp, and we used 6 frequency channels per syllable, the spectrotemporal patterns are matrices with 6 rows and 8 columns for each syllable (ω) (Figure S1).

Because the vector *I* determines the global attractor, sequential activation of the gamma units makes the global attractor change continuously over time and generate the pattern corresponding to syllable ‘*ω*’ when *v^(1)^_ω_ = 1* and *v^(1)^_not ω_* = 0.

The outputs of this level are the state of the Hopfield network, which predicts the activity of the frequency channels in the input, and the causal state associated with the envelope (relayed from the top level):

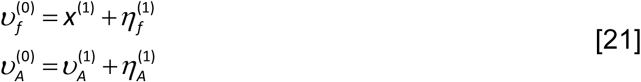

Those quantities are compared with the envelope (*E(t)*) and frequency channels (*F_f_(t)*) in the input signal:

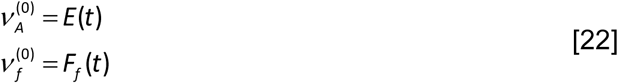

The discrepancy between top-down predictions and sensory input is propagated back in the hierarchy to update hidden state and causal state estimates so that prediction errors are minimized.

The values of all parameters used in the model, as well as precisions for hidden and causal states for both levels, are presented in tables 2 and 3 respectively.

**Table 2.** Parameter values

|  | Hidden states | Causal states |
| --- | --- | --- |
| Top level | $\tau = 10$<br>$b = 0.5, c = 0.04$<br>$\Omega = 4$<br>$\lambda = 0.125, \kappa_2 = 0.2625$<br>$\beta = 3$ | $\alpha = 3$ |
| *Triggers* | $\sigma_1 = 0.005, \sigma_2 = 0.075$<br>$\alpha_1 = 730$ | |
| Bottom level | $\kappa_1 = 2$ | |

**Table 3.** Precisions

|  | Hidden states | Causal states |
| --- | --- | --- |
| Top level | $W_\gamma = \exp(15)$<br>$W_\omega = \exp(2)$<br>$W_\theta = \exp(30)$<br>$W_A = \exp(30)$<br>$W_R = \exp(10)$<br>$W_r = \exp(15), r = (r_1, r_2, r_{2,1}, r_{2,2})$ | $V_\gamma = \exp(5)$<br>$V_\omega = \exp(5)$<br>$V_A = \exp(30)$ |
| Bottom level | $W_f = \exp(8)$ | $V_f = \exp(12)$<br>$V_A = \exp(15)$ |

### Resets/Triggers and Model Variants

To ensure that predictions are temporally aligned with the input, the model needs to align the gamma network with syllable onsets. Moreover, ideally, evidence accumulation should be reset before or at syllable onset. In principle, both resets could be driven by prediction errors. However, our basic model also involves explicit resets.

When present, the trigger to reset gamma units (denoted by T_γ_ in equations 11 and 12) was driven either by local minima in the envelope, which we refer as local minima triggers *T_lm_*, or by theta-filtered local minima (referred as theta triggers *T_θ_*).

**Local minima triggers** *T_lm_* were defined as the periods for which r_2_ ≈ 0 and r_2,2_ > 0 (equations 8 and 9 and red trace in Figure 2B). These were defined as the product of a Gaussian on r_2_ centered at zero and a sigmoid function on r_2,2_:

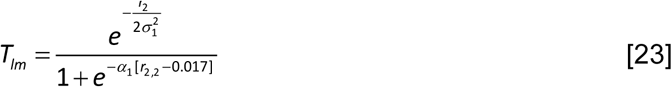

We used a fixed positive threshold 0.017 to reduce false positives due to fluctuations on the envelope.

**Theta-triggers** *T*_θ_: Not all local minima correspond to syllable onsets and the theta oscillation was used to select local minima that are separated by at least one theta cycle. Operationally, this was done by selecting only local minima that occurred in the vicinity of the theta oscillation minima, defined by phase eligibility windows *T*φ** (red trace in Figure 2A)

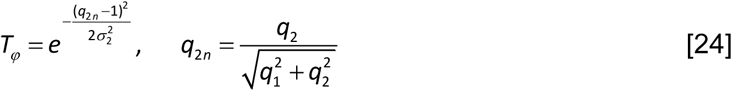

The overlap between the local minimum trigger *T_lm_* and the eligibility window *T*φ**, defines the theta triggers *T*_θ_ (red trace in Figure 2C):

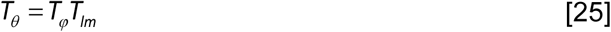

When present, the reset to syllable units (*Tω*, equation 14) was driven by the theta triggers *T_θ_* (defined above) or by the model’s knowledge about syllable duration provided by the sequence of gamma units. Since the last (eighth) gamma unit signals the end of the syllable, it can define a trigger that we refer as the internal trigger (denoted as *T_int_* and plotted as a thick line in figure 2E):

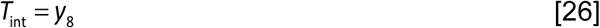

In summary, to reset gamma units the model uses envelope based estimates of syllable onset locations provided either by local minima or theta triggers (*T_lm_* and *T_θ_* respectively), whereas to reset syllable units the model either uses theta triggers (*T_θ_*) or internal information about syllable duration -*T_int_*. To explore the relative importance of each resetting mechanism for the overall performance of the model, we compared five different model variants (Figure 3, top panel); each with a different combination of triggers to reset the dynamics of syllable and gamma units (Table 1).

### Model Output

The performance of the model was quantified by the proportion of correctly identified syllables. To define the syllables identified by the model, we considered the time average of the causal state (ω, equation 14) of each of the syllable units taken within the boundaries of each input syllable. The syllable unit with the highest causal state average over this window was considered the model’s choice.

The temporal window is determined by the actual input syllable boundaries and not by the model’s estimates of syllable boundaries, since the latter changed across models.

When the model’s choice corresponded to the syllable in the input, we considered it as successful recognition. For each sentence, we calculated the percentage of correctly identified syllables.

### Statistical analyses

As described in the previous section, a single number (% correctly identified syllables) describes the performance of the model for each sentence. Simulations were run on 30 sentences, and the performance of each model architecture is thus described by a vector of 30 numbers. The non-parametric Wilcoxon signed-rank test for repeated measures was used to compare models’ performance. False discovery rate (Benjamini and Hochberg, 1995; Benjamini and Yekutieli, 2001) was used to control for multiple comparisons.

Finally, chance level (around 8%) for the whole dataset was estimated by calculating the median of the distribution of the chance level across all sentences. The chance level of each sentence equals to 1/N*_syl.,_* where N*_syl_* is the number of syllables in it.

