## Supplementary material for "Combining predictive coding with neural oscillations optimizes on-line speech processing"

### SUPPLEMENTAL INFORMATION

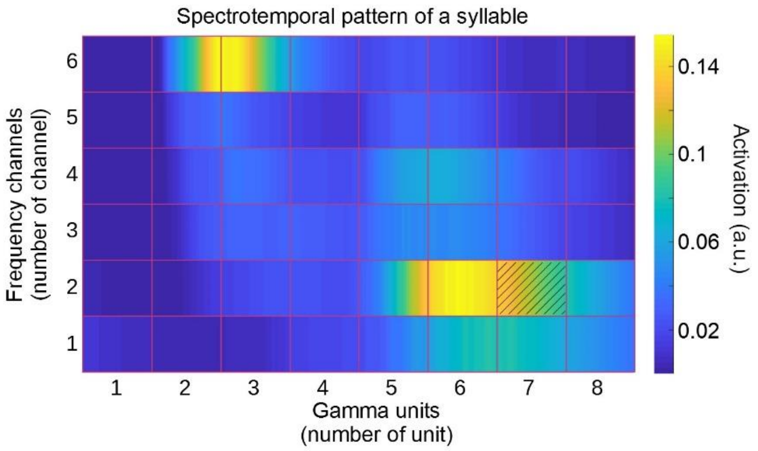


**Figure S1. Extraction of spectrotemporal patterns, related to Figure 1.** The figure illustrates how the spectrotemporal pattern ($ST_{f\gamma\omega}$) of each syllable $\omega$ was calculated. As we have six frequency channels and eight gamma units per syllable, we created 6x8 matrices where each entry corresponds to the average amplitude of the associated frequency channel over the duration of a given gamma unit. For example, the entry $ST_{27\omega}$ (f=2 and $\gamma$=7), dashed entry on the figure, represents the average activation of the second frequency band within the seventh gamma cycle (equation 19).

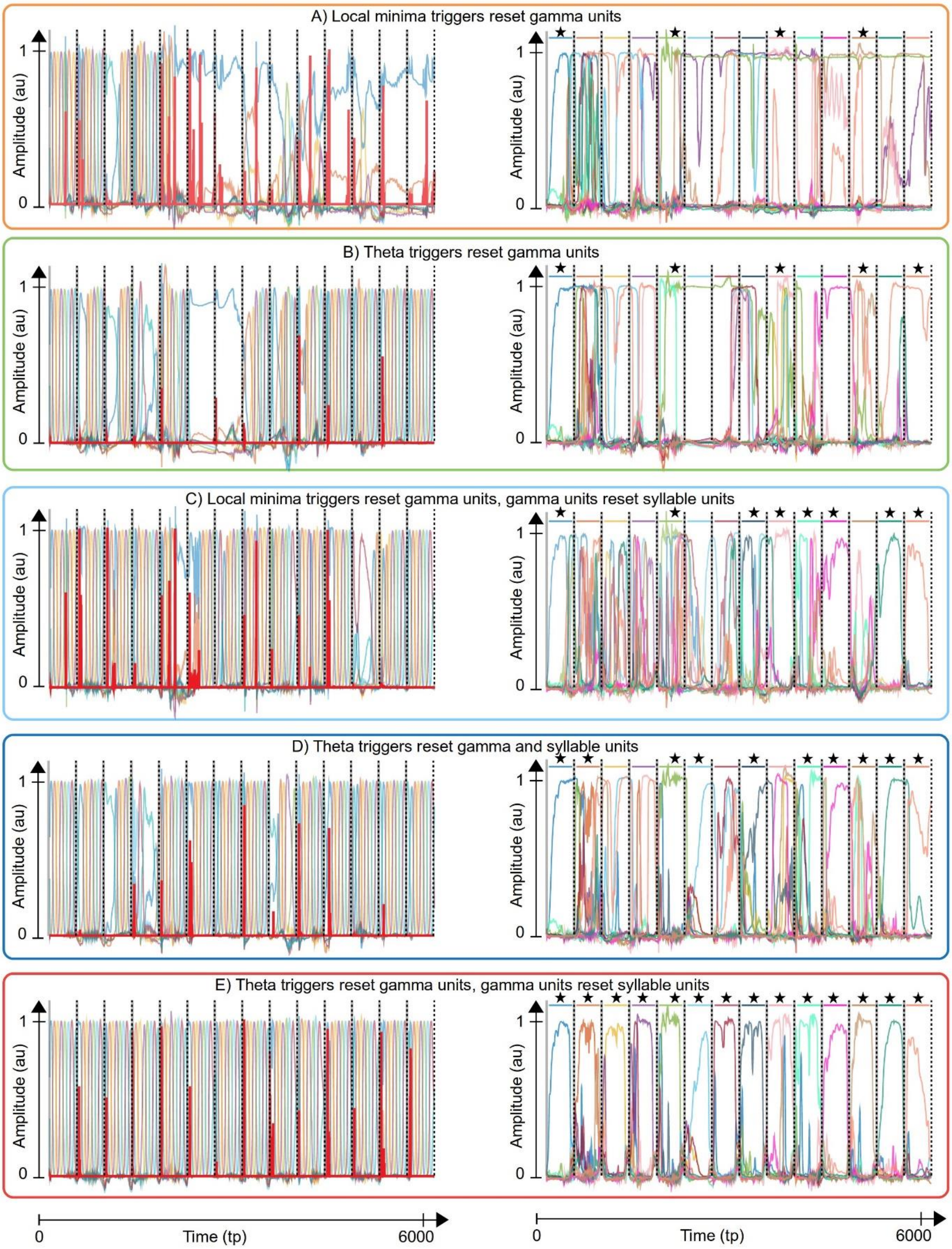


**Figure S2. Dynamics of syllable and gamma units, related to Figure 2.** Activity of gamma (left column) and syllable units (right column) for the 5 model configurations shown in Figure 3. Simulations correspond to the same sentence from the same speaker. For reference, all panels show the location of the veridical syllable onsets (vertical grey lines) and offsets (vertical black dash lines). The triggers that activate the 1$^{st}$ gamma unit are shown in thick red lines, while gamma units are shown in different colours with the 1$^{st}$ gamma unit, which initiates the gamma sequence, drawn in blue. Syllable units, on the right column, are shown in different colours. There are as many syllable units as syllables in the input sentence, and the bars at the top indicate the colour corresponding to the veridical syllable in the input at each syllabic interval. A star above a bar indicates that the model selected the right syllable (the selected syllable was defined as the syllabic unit with highest average activation within the syllabic interval). A) Gamma units are reset by local minima in the envelope and syllable units are not reset. Resulting gamma activity was distorted (note the breakdown of their sequential activation) and syllable units failed to identify clear winners within syllable boundaries. B) Enabling theta-gamma coupling (reset of gamma units by theta detected onsets) results in a less distorted gamma sequence (breakdown of their sequential activation occurred less often). However syllable recognition did not improve, likely because syllable units were not reset. C) Enabling the reset of syllable units based on endogenous syllable duration information improved the number of correctly selected syllables (note the increase in overall performance in Figure 3). This happened despite gamma units being reset (as in A) by local minima triggers; as syllable units were temporally organized, there were clearer peaks for recognized syllables (marked with stars), leading to a better-organized gamma sequence (only very short periods with breakdown of sequential activation). D) Theta-gamma coupling is combined with a theta-driven reset of syllable units. Despite the theta-gamma coupling, there was no clear improvement in performance with respect to model C. E) Theta-gamma coupling is now combined with reset of syllable units by the last gamma unit. There was no breakdown of sequential activation of gamma units and syllable units now showed clear peaks within syllable boundaries, which for the present example resulted in the correct recognition of all syllables.

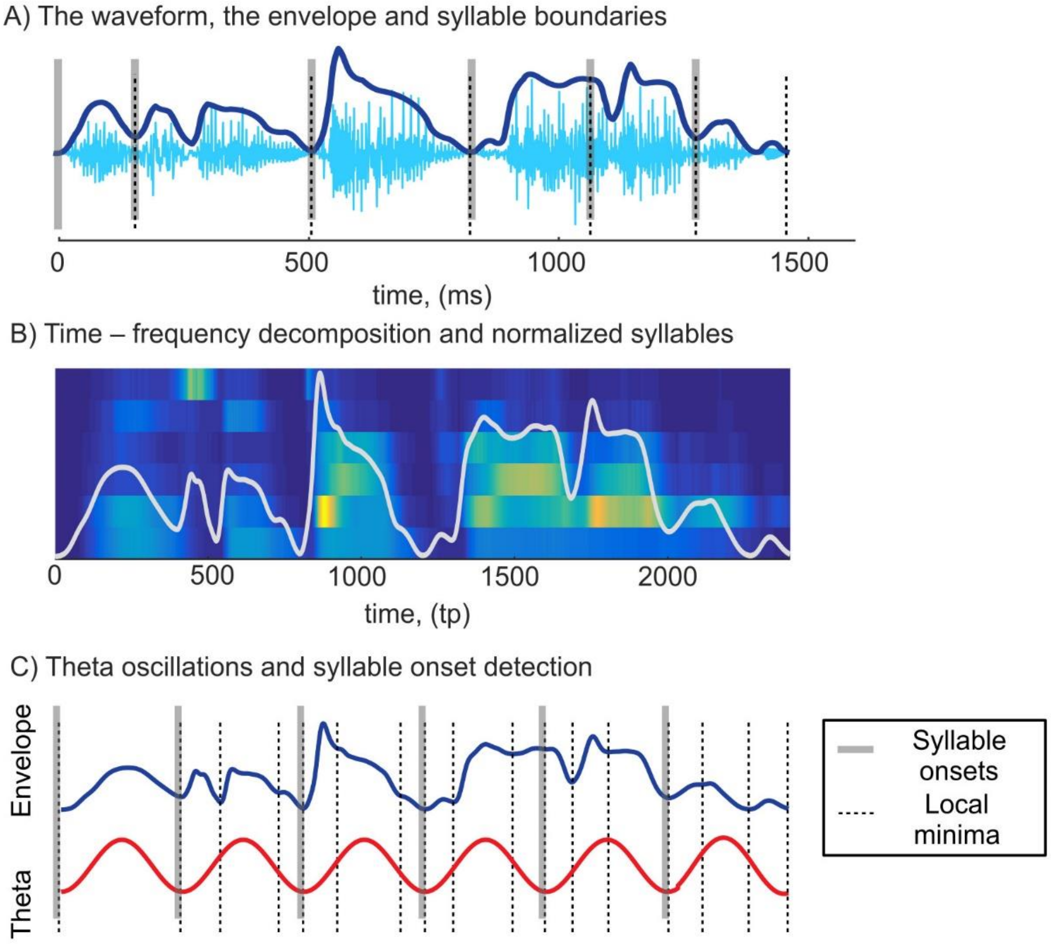


**Figure S3. Preprocessing of the input, related to Figure 1.** Simulations were run with natural sentences from the TIMIT database. A) Waveform, envelope and syllable boundaries of the sentence "They seem darned proud of it" B) Pre-processed sentence used for simulations. We used a biologically inspired model of auditory periphery to calculate the auditory spectrogram and reduced to 6 frequency channels, normalized to obtain a mean syllable duration of 200 ms, and resampled to 400-time points (samples). Possible gaps between syllables were also resampled. C) Syllable onsets are located next to local minima in the envelope (thin dashed lines), i.e. a low energy articulatory footprint corresponding to lip closure between words/syllable. Importantly, as not every local minimum signals syllable onset, we hypothesize that the brain uses theta oscillations to select only local minima that are separated by at least one theta cycle.

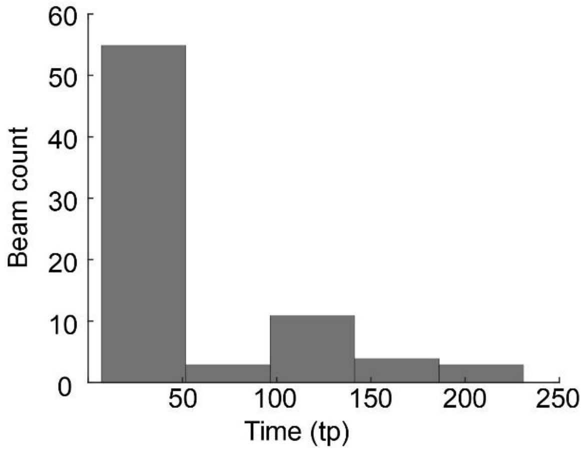


**Figure S4. Gap distribution in input sentences, related to Figure 1.** Distribution of the gaps in the 30 sentences used for simulations. Overall, there are 76 gaps, most of which have duration within 1-25 ms (on the figure gap duration is represented as time points (tp) (with a 1 ms ≈ 2 tp, sampling rate).

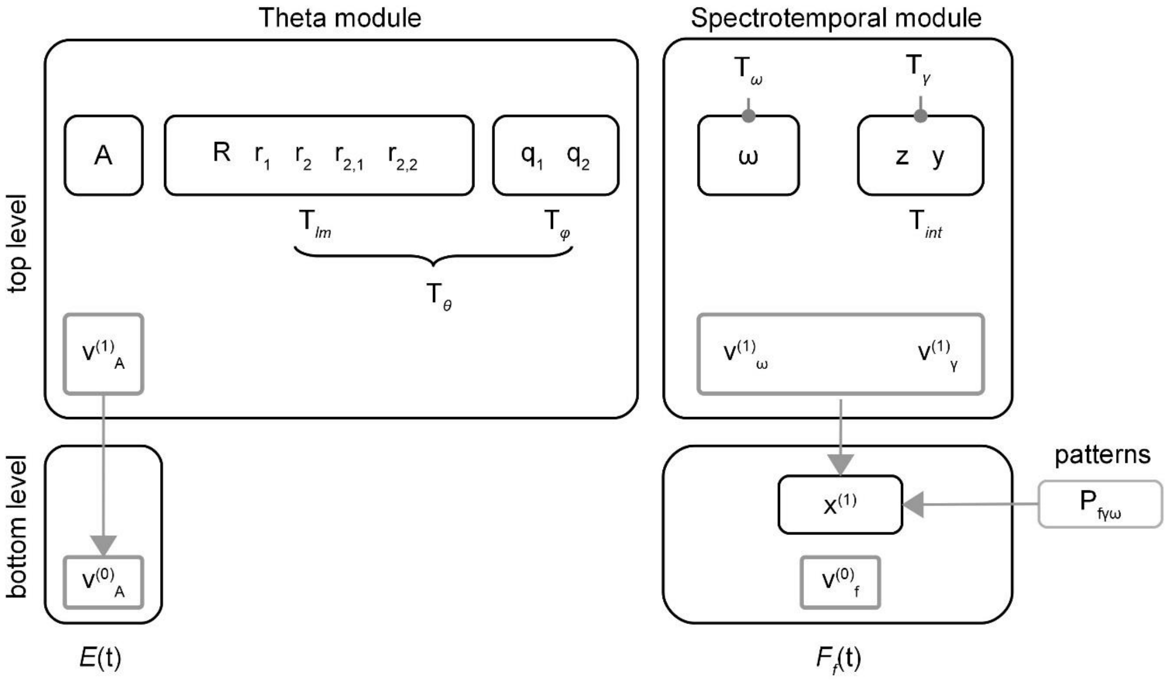


**Figure S5. Schematics of the model, related to Figure 2:** The figure shows all the variables in the model. Black boxes include the hidden states of each level, whereas gray boxes correspond the information that each level passes to the level below. The top level sends information about the envelope $v_A^{(1)}$ and the dynamics of the syllable $v_\omega^{(1)}$ and gamma units $v_\gamma^{(1)}$. The output of the first level ($v_A^{(0)}$ for the envelope and $v_f^{(0)}$ for the frequency channels) is then compared with the input signal (E(t) and $F_f$(t) respectively).

**Table S1. List of sentences used for simulations, related to Figure 3.** The three speakers used for simulations come from the same dialect (DR8) in the TIMIT database. The first column indicates sentence number and corresponding ID in the database. The middle column shows speaker ID, e.g. for Speaker 3 MMWS0, where the first letter corresponds to gender. The last column shows the number of syllables in the sentence. Overall, for Speaker 1 we have 131 syllables, for the second and third speakers 138 and 120 syllables respectively. In total, 389 syllables were used.

| Sent. ID | Sentences, speaker 1, TIMIT ID = MBCG0, dialect=DR8 | N syll. |
|---|---|---|
| 1.SA1 | She had your dark suit in greasy wash water all year. | 14 |
| 2.SA2 | Don’t ask me to carry an oily rag like that. | 13 |
| 3.SI486 | It was time to go up myself. | 8 |
| 4.SI957 | But he was very much like his hatred of camp routine. | 19 |
| 5.SI2217 | He saw the dangers, not the glories of being identified as a mutineer. | 11 |
| 6. SX57 | The prowler wore a ski mask for disguise. | 10 |
| 7.SX147 | Correct execution of my instructions is crucial. | 13 |
| 8. SX237 | The courier was a dwarf. | 8 |
| 9.SX327 | Al received a joint appointment in the biology and the engineering departments. | 23 |
| 10.SX417 | Valley Lodge yearly celebrates the first calf born. | 12 |
| | **Sentences, speaker 2, TIMIT ID = MRLK0, dialect=DR8** | |
| 1.SA1 | She had your dark suit in greasy wash water all year. | 14 |
| 2.SA2 | Don’t ask me to carry an oily rag like that. | 12 |
| 3.SI843 | To begin with what is an interior designer.8 | 15 |
| 4.SI1468 | A precision transit is set up so that it is lined with respect to true north. | 21 |
| 5.SI2140 | Was it a hysterical release from the long strain of vigilance of those weeks? | 21 |
| 6.SX33 | Coconut cream pie makes a nice dessert. | 10 |
| 7.SX123 | A screwdriver is made from vodka and orange juice. | 13 |
| 8.SX213 | The news agency hired a great journalist. | 12 |
| 9.SX303 | The bluejay flew over the high building. | 10 |
| 10.SX493 | She uses both names interchangeably. | 10 |
| | **Sentences, speaker 3, TIMIT ID = MMWS0, dialect=DR8** | |
| 1.SA1 | She had your dark suit in greasy wash water all year. | 14 |
| 2.SA2 | Don’t ask me to carry an oily rag like that. | 12 |
| 3.SI559 | They seem darned proud of it. | 6 |
| 4.SI888 | Is a dream simply a mental or cerebral movie? | 14 |
| 5.SI1518 | His arm moved swiftly violently once twice. | 11 |
| 6.SX78 | Doctors prescribe drugs too freely. | 8 |
| 7.SX168 | Who took the kayak down the bayou? | 9 |
| 8.SX258 | The essay undeniably reflects our view ably. | 15 |
| 9.SX348 | I'll have a scoop of that exotic purple and turquoise sherbet. | 16 |
| 10.SX438 | Shell shock caused by shrapnel is sometimes cured through group therapy? | 15 |

**Table S2. Corrected p-values for pairwise comparisons, related to Figure 3.** Corrected *p*-values for pairwise comparisons. Significant level p<0.0005. Blue cells correspond to significant performance difference within a selected pair, whereas red cells correspond to conditions in which the difference is not statistically significant.

| | Model A) | Model B) | Model C) | Model D) | Model E) |
|---|---|---|---|---|---|
| Model A) | | 0.186 | 9.32e-6 | 6.38e-6 | 6.38e-6 |
| Model B) | 0.186 | | 9.32e-6 | 9.32e-6 | 6.38e-6 |
| Model C) | 9.32e-6 | 9.32e-6 | | 0.681 | 1.50e-4 |
| Model D) | 6.38e-6 | 9.32e-5 | 0.681 | | 1.26e-4 |
| Model E) | 6.38e-6 | 6.38e-6 | 1.50e-4 | 1.26e-4 | |
